# Rapid and sensitive virulence prediction and identification of Newcastle disease virus genotypes using third-generation sequencing

**DOI:** 10.1101/349159

**Authors:** Salman Latif Butt, Tonya L. Taylor, Jeremy D. Volkening, Kiril M. Dimitrov, Dawn Williams-Coplin, Kevin K. Lahmers, Asif Masood Rana, David L. Suarez, Claudio L. Afonso, Adrian Marchetti

## Abstract

Newcastle disease (ND) outbreaks are global challenges to the poultry industry. Effective management requires rapid identification and virulence prediction of the circulating Newcastle disease viruses (NDV), the causative agent of ND. However, these diagnostics are hindered by the genetic diversity and rapid evolution of NDVs. A highly sensitive amplicon sequencing (AmpSeq) workflow for virulence and genotype prediction of NDV samples using a third-generation, real-time DNA sequencing platform is described using both egg-propagated virus and clinical samples. 1D MinION sequencing of barcoded NDV amplicons was performed on 33 egg-grown isolates, (23 unique lineages, including 15 different NDV genotypes), and from 15 clinical swab samples from field outbreaks. Assembly-based data analysis was performed in a customized, Galaxy-based AmpSeq workflow. For all egg-grown samples, NDV was detected and virulence and genotype were predicted. For clinical samples, NDV was detected in ten of eleven NDV samples. Six of the clinical samples contained two mixed genotypes, of which the MinION method detected both genotypes in four of those samples. Additionally, testing a dilution series of one NDV sample resulted in detection of NDV with a 50% egg infectious dose (EID50) as low as 10^1^ EID_50_/ml. This was accomplished in as little as 7 minutes of sequencing time, with a 98.37% sequence identity compared to the expected consensus. The high sensitivity, fast sequencing capabilities, accuracy of the consensus sequences, and the low cost of multiplexing allowed for identification of NDV of different genotypes circulating worldwide. This general method will likely be applicable to other infections agents.

## Introduction

Newcastle disease (ND) is one of the most important infectious diseases of poultry and is a major cause of economic losses to the global poultry industry. Newcastle disease is caused by virulent strains of avian paramyxovirus 1 (APMV-1), commonly known as Newcastle disease virus (NDV) (1), recently reclassified as avian avulavirus-1 (AAvV-1) (2). Newcastle disease viruses are a highly diverse group of viruses with two distinct classes and 19 accepted genotypes, infecting a wide range of domestic and wild bird species. In addition to the genotypic diversity of NDVs, these viruses are also diverse in their virulence. The global spread, constant evolution, varying virulence, and the wide host range of NDV are challenges to the control of ND (3).

Effective control of ND is dependent on rapid, sensitive, and specific diagnostic testing, which for ND are typically oriented towards detection, genotyping, or prediction of virulence. Virulence of NDV is best assayed through infection-based studies (4), but due to the time constraints associated with such methods, reverse transcriptase-quantitative PCR (RT-qPCR) and sequencing of the fusion (F) gene cleavage site are used to predict NDV virulence (5, 6). Genotyping of NDV is commonly achieved through Sanger sequencing of the coding sequence of the fusion gene (7), which also allows for prediction of virulence. Preliminary genotyping can be accomplished through partial fusion gene sequencing (i.e., variable region) (8). PCR-based tests aimed at rapid detection often lack applicability for virulence determination due to NDV’s genetic diversity, and the current methods that rely on Sanger sequencing lack multiplexing capability and have limited sequencing depth, which complicates detection of mixed infections. While fusion-based assays can be used for detection (9), the variability of this region, which makes it useful for genotyping, hinders the universal applicability of any single primer set. Thus, detection-focused assays are often designed towards more conserved regions, such as the matrix or polymerase genes(9–11). These assays, however, fail to provide virulence or genotype predictions. In summary, there is a need for a method that will sensitively and rapidly detect numerous genotypes of NDV and provide genotype and virulence prediction.

Rapid advances in nucleic acid sequencing, have led to different sequencing platforms (12, 13) being widely applied for identification of novel viruses (14), whole genome sequencing (15), transcriptomics, and metagenomics (16, 17). However, high capital investments and relatively long turnaround times limit the widespread use of these NGS platforms, especially in developing countries (18). Recent improvements in third-generation sequencing, including those introduced by Oxford Nanopore Technologies (ONT) (19), increase the utility of high-throughput sequencing as a useful tool for surveillance and pathogen characterization (20). Among the transformative advantages of ONT’s sequencing technology are the ability to perform real-time sequence analysis with a short turnaround time (21), the portability of the MinION device, the low startup cost compared to other high-throughput platforms, and the ability to sequence up to several thousand bases from individual RNA or DNA molecules. The MinION device has been successfully used to evaluate antibiotic resistance genes from several bacterial species (22, 23), obtain complete viral genome sequences of an influenza virus (24) and Ebola virus (25), and detect partial viral genome sequences (e.g., Zika virus (26) and poxviruses (21)) by sequencing PCR amplicons (AmpSeq). The MinION, therefore, represents an opportunity to take infectious disease diagnostics a step further and to perform rapid identification and genetic characterization of infectious agents at a lower cost.

As with any deep sequencing platform, the sequence analysis approach is integral for accurate interpretation. Primarily, two approaches for taxonomic profiling of microbial sequencing data have been employed: read-based and *de novo* assembly-based classifications. Read-based metagenomic classification software has been used for identification of microbial species from high-throughput sequencing data (19, 27–29). Although the sequencing accuracy of the MinION is improving, the raw single-read error rate of nearly 10% (30) may limit the accuracy of this approach for Nanopore data (27), especially when attempting to subspecies level differentiation. *De novo* approaches that use quality-based filtering and clustering of reads (31), or use consensus-based error correction of Nanopore sequencing reads have been reported (32); however, these are not optimized for amplicon sequencing data.

In this study, a specific, sensitive, rapid protocol, using the MinION sequencer, was developed to detect representative isolates from all current (excluding the Madagascar-limited genotype XI) genotypes of NDV. This protocol was also tested on a limited number of clinical swab samples collected from chickens during disease outbreaks. Additionally, a Galaxy-based, *de novo* AmpSeq workflow is presented that efficiently reduces systematic sequencing errors in Nanopore sequencing data and uses amplicon-based sequences to obtain accurate final consensus sequences.

## Materials and Methods

### Viruses

Thirty-three NDV isolates, representing 23 unique lineages (including 15 different genotypes) of different virulence, and 10 other avian avulaviruses (AAvV 2-10 and AAvV-13) from the Southeast Poultry Research Laboratory (SEPRL) repository, were propagated in 9–11-day-old specific pathogen free (SPF) eggs (33) and the harvested allantoic fluids were used in this study. Additionally, 15 oral and cloacal swab samples collected from chickens during disease outbreaks in Pakistan in 2015 were tested. The background information of the egg-grown isolates and the clinical samples is summarized in Table S1 and Table S2, respectively.

### RNA Isolation

Total RNA from each sample was extracted from infectious allantoic fluids or directly from clinical swab media using TRIzol LS (Thermo Fisher Scientific, USA) following the manufacturer’s instructions. RNA concentrations were determined by using Qubit® RNA HS Assay Kit on a Qubit® fluorometer 3.0 (Thermo Fisher Scientific, USA).

### Amplicon synthesis

Approximately 20 ng (in 5 μl) of RNA was reverse transcribed, and cDNA was amplified with target-specific primers using the SuperScript™ III One-Step RT-PCR System (Thermo Fisher Scientific, USA). Previously published primers (4331F and 5090R) (8, 34) were used in this protocol; however, primers were tailed with universal adapter sequence of 22 nucleotides (in bold font) to facilitate PCR-based barcoding: 4331F Tailed: 5′-**TTTCTGTTGGTGCTGATATTGC**GAGGTTACCTCYACYAAGCTRGAGA-3′; 5090R Tailed: 5′-**ACTTGCCTGTCGCTCTATCTTC**TCATTAACAAAYTGCTGCATCTTCCCWAC-3′). The thermocycler conditions for the reaction were as follows: 50 °C for 30 minutes; 94 °C for 2 minutes; 40 cycles of 94 °C for 15 seconds, 56 °C for 30 seconds, and 68 °C for 60 seconds, followed by 68 °C for 5 minutes. The reaction amplified a 788 base pair (bp) NDV product (832 bp including primer tails) for all genotypes, which included 173 bp of the 3′ region of the end of the M gene and 615 bp of the 5′ end of the F gene (sizes and primer locations based on the Genotype V strain).

### Library preparation

Amplified DNA was purified by Agencourt AMPure XP beads (Beckman Coulter, USA) at 1.6:1 bead-to-DNA ratio and quantified using the dsDNA High Sensitivity Assay kit on a Qubit® fluorometer 3.0 (Thermo Fisher Scientific, USA). MinION-compatible DNA libraries were prepared with approximately 1 μg of barcoded DNA in a total volume of 45 μL using nuclease-free water and using the 1D PCR Barcoding Amplicon Kit (Oxford Nanopore Technologies, UK) in conjunction with the Ligation Sequencing Kit 1D (SQK-LSK108) (19) as per manufacturer’s instructions. Briefly, each of the amplicons were diluted to 0.5 nM for barcoding and amplified using LongAmp Taq 2X Master Mix (New England Biolabs, USA) with the following conditions; 95 °C for 3 min; 15 cycles of 95 °C for 15 seconds; 62 °C for 15 seconds, 65 °C for 50 seconds, followed by 65 °C for 50 seconds. The barcoded amplicons were bead purified, pooled into a single tube, end prepped, dA tailed, bead purified, and ligated to the sequencing adapters per manufacturer’s instructions. Final DNA libraries were bead purified and stored frozen until used for sequencing.

### Comparison to RT-qPCR assay for Matrix gene

For comparison of this MinION-based protocol with the matrix gene reverse transcriptase-quantitative polymerase chain reaction (RT-qPCR) assay, both methods were run on a dilution series from a single isolate. NDV (LaSota strain) from the SEPRL repository was cultured in SPF 9–11-days-old eggs and the harvested allantoic fluids were diluted to titers ranging from 10^6^ to 10^1^ EID_50_/mL in brain-heart infusion broth. RNA was extracted from dilutions, and DNA libraries were prepared following the same protocols as described above. Amplicons from each of the dilutions were barcoded separately. At the pooling step, equal concentrations of barcoded amplicons from different dilutions of LaSota were pooled together in single tube. Dilutions, extractions, library construction, and sequencing were performed twice.

The same extracted RNA was also used as the input into the RT-qPCR using the AgPath-ID one-step RT-PCR Kit (Ambion, USA) on the ABI 7500 Fast Real-Time PCR system following the previously described protocols (9).

### Sequencing by MinION

The libraries were sequenced with the MinION Nanopore sequencer (19). A new FLO-MIN106 R9.4 flow cell, stored at 4°C prior to use, was allowed to equilibrate to room temperature for 10 minutes before priming it for sequencing. The flow cell was primed with running buffer as per manufacturer’s instructions. The pooled DNA libraries were prepared by combining 12 μL of the libraries with 2.5 μL nuclease-free water, 35 μL RBF, and 25.5 μL library loading beads. After the MinION Platform QC run, the DNA library was loaded into the MinION flow cell via the SpotON port. The standard 48-h 1D sequencing protocol was initiated using the MinKNOW software v.5.12. Detailed information for all MinION runs in this study is provided in Table S3.

The complete steps from RNA isolation to MinION sequencing were performed twice for egg-grown viruses. One run consisted of six egg-grown isolates from different genotypes representative of vaccine and virulent NDV strains (run 3: 6-sample pool). The other run consisted of these same six viruses and an additional 27 egg-grown NDV isolates (run 4: 33-sample pool).

To determine the accuracy of consensus sequences at different sequencing time points for accurate identification of the NDV genotypes, the raw data (FAST5 files) obtained from the 10-fold serial dilution experiment (see above) were analyzed in subgroups based on time of acquisition and processed through the AmpSeq workflow as described below.

### MinION data analysis workflow

To analyze the Nanopore sequencing data, a custom, assembly-based AmpSeq workflow within the Galaxy platform interface (35) was developed, as diagrammed in Figure 1. The MinION raw reads in FAST5 format were uploaded into Galaxy workflow. The reads were base-called using the Albacore v2.02 (ONT). The NanoporeQC tool v0.001 (available in the Galaxy testing toolshed) was used to visualize read quality based on the summary table produced by Albacore. Porechop v0.2.2 (https://github.com/rrwick/Porechop) was used to demultiplex reads for each of the barcodes and trim the adapters at the ends of the reads by using default settings. Short reads (cutoff = 600 bp) were filtered out and the remaining reads were used as input to the in-house LAclust v0.002. LAclust performs single-linkage clustering of noisy reads based on alignment identity and length cutoffs from DALIGNER pairwise alignments (36) (minimum alignment coverage = 0.90, maximum identity difference = 0.35; minimum number of reads to save cluster = 5; maximum reads saved per cluster = 200, minimum read length = 600 bp; rank mode = number of intracluster linkages; randomized input read order = yes). Read clusters generated by LAclust were then aligned using the in-house Amplicon aligner v0.001 to generate a consensus sequence. This tool optionally subsamples reads (target depth used = 100), re-orients them as necessary, aligns them using Multiple Alignment using Fast Fourier Transform (MAFFT) (37) with highly relaxed gap opening and extension penalties, and calls a majority consensus. Next, each consensus was used as a reference sequence for mapping the full unfiltered read clusters from LAclust with BWA-MEM and ONT2D settings (38, 39). The final consensus sequence for each sample was refined by using Nanopolish v0.8.5 (https://github.com/jts/nanopolish), which calculates an improved consensus using the read alignments and raw signal information from the original FAST5 files. After manually trimming primer sequences from both 3′ (25 bp) and 5′ (29 bp) ends, the obtained consensus sequences (734 bp) were BLAST searched against NDV customized database, which consisted NCBI’s nucleotide (nt) database and internal unpublished NDV sequences (NCBI database updated on May 23, 2018).

**Figure 1.**
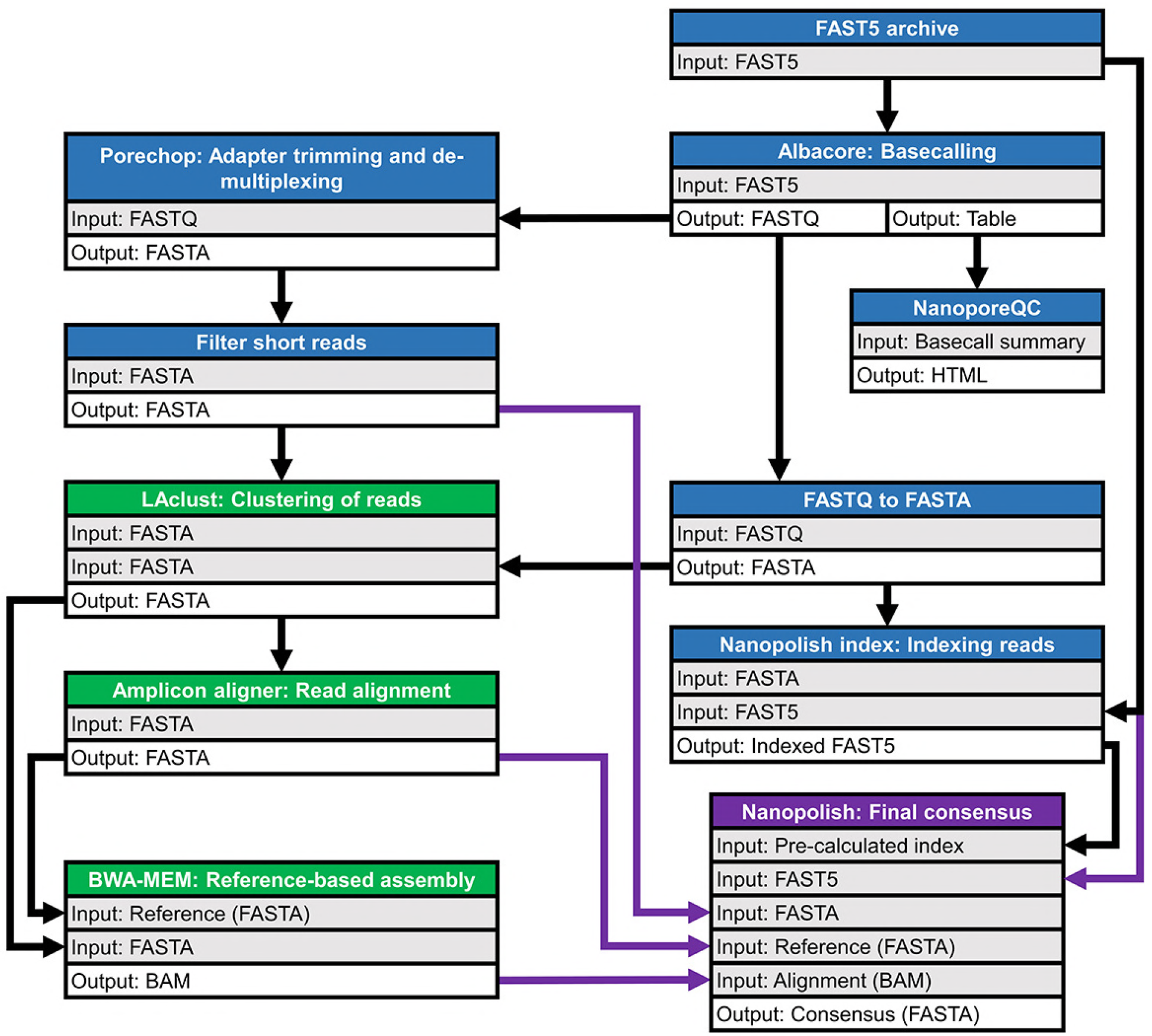
Schematic diagram of customized Galaxy workflow for MinION sequence data analysis. *Blue shading* indicates pre-processing steps. *Green shading* indicates post-processing steps; assembly/output is shaded purple. *Purple* arrows indicate different inputs for final consensus calculation.

### Sequencing by MiSeq

For comparison between nucleotide sequences obtained from MinION and MiSeq (a validated and high accuracy sequencing platform), 24 NDV isolates from the SEPRL repository that were used for MinION sequencing (representing each genotype and sub-genotype of NDV) and 15 clinical swab samples (allantoic fluid of cultured swab samples) were processed for target-independent NGS sequencing. Briefly, paired-end random sequencing was conducted from cDNA libraries prepared from total RNA using KAPA Stranded RNA-Seq kit (KAPA Biosystems, USA) as per manufacturer’s instructions and as previously described (40). All libraries for NGS were loaded into the 300-cycle MiSeq Reagent Kit v2 (Illumina, USA) and pair-end sequencing (2 × 150 bp) was performed on the Illumina MiSeq instrument (Illumina, USA). Pre-processing and de-novo assembly of the raw sequencing data was completed within the Galaxy platform using a previously described approach (15).

### Phylogenetic analysis

The assembled consensus sequences from different NDV genotypes and sub-genotypes (6 sequences from MinION run 3, 32 sequences from run 4 and 24 sequences from MiSeq; a total of 62 sequences) and selected (minimum of one sequence from each genotype/subgenotype) sequences from GenBank (n = 66) were aligned using ClustalW (41) in MEGA6 (42). Determination of the best-fit substitution model was performed using MEGA6, and the goodness-of-fit for each model was measured by corrected Akaike information criterion (AICc) and Bayesian information criterion (BIC) (42). The final tree was constructed using the maximum-likelihood method based on the General Time Reversible model as implemented in MEGA6, with 500 bootstrap replicates (43). The available GenBank accession number for each sequence in the phylogenetic tree is followed by the, host name, country of isolation, strain designation, and year of isolation.

### Comparison of MinION and MiSeq sequence accuracy

To assess the accuracy of the MinION AmpSeq consensus sequences, 24 samples were sequenced by both deep-sequencing methods (MinION and MiSeq) described above. Pairwise nucleotide comparison between MinION and MiSeq was conducted using the Maximum Composite Likelihood model (44). The variation rate among sites was modeled with a gamma distribution (shape parameter = 1). The analysis involved 54 nucleotide sequences. Codon positions included were 1st+2nd+3rd+Noncoding. All positions containing gaps and missing data were eliminated. There were a total of 691 positions in the final dataset. The evolutionary distances were inferred by pairwise analysis using the MEGA6 (42).

### Accession number

The sequences obtained in the current study were submitted to GenBank and are available under the accession numbers from MH392212 to MH392228.

## Results

### Comparison to the Matrix gene RT-qPCR assay

Six, sequential, 10-fold dilutions (from 10^6^ EID_50_/ml to 10^1^ EID_50_/ml) from one NDV isolate (LaSota) were used to compare the ability of AmpSeq and RT-qPCR to detect low quantities of NDV. In each of the six dilutions, AmpSeq and the matrix RT-qPCR detected NDV in all dilutions. AmpSeq resulted in 99.04–100.0% sequence identity to the LaSota isolate across all six dilutions in the first experiment (R1) and 99.86–100.0% identity in the second experiment (R2) (Table 1).

**Table 1.** Comparison of MinION sequencing to RT-qPCR 1 for detection of NDV LaSota

| Dilution<br>(EID50/ml) | Total reads | Total NDV<br>reads | Reads per<br>consensus<sup>c</sup> | Percent<br>identity<sup>d</sup> | Consensus<br>length | RT-qPCR<sup>e</sup><br>(Ct) |
| --- | --- | --- | --- | --- | --- | --- |
|  | R1<sup>a</sup> \| R2<sup>b</sup> | R1 \| R2 | R1 \| R2 | R1 \| R2 | R1 \| R2 | R1 \| R2 |
| 10<sup>6</sup> | 6667 \| 11366 | 6577 \| 10861 | 200 \| 200 | 100 \| 100 | 734 \| 734 | 21.8, 21.1 \| 22.7, 22.7 |
| 10<sup>5</sup> | 4519 \| 6801 | 4439 \| 6540 | 200 \| 200 | 100 \| 100 | 734 \| 734 | 26.3, 25.8 \| 26.2, 26.4 |
| 10<sup>4</sup> | 3856 \| 8289 | 3829 \| 7890 | 200 \| 200 | 100 \| 100 | 734 \| 734 | 28.9, 27.8 \| 29.1, 29.3 |
| 10<sup>3</sup> | 164 \| 9484 | 157 \| 9061 | 157 \| 200 | 100 \| 100 | 734 \| 734 | 31.4, 31.1 \| 32.7, 32.8 |
| 10<sup>2</sup> | 94 \| 4939 | 85 \| 4725 | 81 \| 200 | 100 \| 100 | 734 \| 734 | 34.2, 34.8 \| 34.8, 35.3 |
| 10<sup>1</sup> | 133 \| 2652 | 131 \| 2520 | 73 \| 200 | 99.04 \| 99.86 | 729 \| 734 | 34.9, 34.8 \| 36.9, 36.7 |

<sup>a</sup>Replicate 1
<sup>b</sup>Replicate 2
<sup>c</sup>*LAclust* was limited (using the available options) to only use 200 reads for downstream analysis
<sup>d</sup> Consensus sequence identity to the reference sequence of NDV LaSota sequenced with MiSeq
(MH392212/chicken/USA/LaSota/1946)
<sup>e</sup>Each dilution performed in duplicate and Ct values from each well are shown here.

**Note:** All 60,000 reads obtained during 32 minutes of sequencing run (R1) were utilized for the analysis. All 98,916 reads from R2 were utilized for the analysis.

### Time for data acquisition and analysis

To determine the minimal sequencing time needed for acquisition of accurate full-length amplicon consensus sequences at different serial dilutions, 28,000 reads, which were obtained within the first 19 minutes of sequencing in the first serial dilution experiment (run 1), were analyzed. For all concentrations, the first read that aligned to the reference LaSota sequence was obtained within 5 minutes after the sequencing run started. To obtain consensus sequences (5 reads required to build a consensus sequence) only 5 minutes of sequencing time were required for concentrations 10^6^–10^3^ EID_50_/ml, which resulted in 99.18–100% sequence identity to the reference LaSota strain. Seven minutes were required to obtain NDV consensus sequences for the two lower concentrations: 10^1^ EID50/ml = 8 reads, 98.77% identity and 10^2^ EID_50_/ml = 5 reads, 98.37% identity (Table 2).

**Table 2.** Accuracy of consensus sequence identity from 10-fold serial dilutions (EID_50_/ml) of NDV LaSota at different time points during MinION sequencing run.

| Sequencing | Total | 10<sup>6</sup> |  | 10<sup>5</sup> |  | 10<sup>4</sup> |  | 10<sup>3</sup> |  | 10<sup>2</sup> |  | 10<sup>1</sup> |  |
| --- | --- | --- | --- | --- | --- | --- | --- | --- | --- | --- | --- | --- | --- |
| run time | Raw | NDV | % | NDV | % | NDV | % | NDV | % | NDV | % | NDV | % |
| (min) | reads | reads<sup>a</sup> | Identity<sup>b</sup> | reads | Identity | reads | Identity | reads | Identity | reads | Identity | reads | Identity |
| 5 | 4000 | 283 | 100 | 199 | 100 | 182 | 100 | 10 | 99.18 | 2 | NA | 3 | NA |
| 7 | 8000 | 644 | 100 | 406 | 100 | 368 | 100 | 14 | 99.32 | 8 | 98.77 | 5 | 98.37 |
| 10 | 12000 | 1029 | 100 | 661 | 100 | 571 | 100 | 23 | 99.32 | 16 | 99.32 | 8 | 99.18 |
| 12 | 16000 | 1397 | 100 | 901 | 100 | 759 | 100 | 29 | 99.86 | 17 | 99.32 | 14 | 99.45 |
| 14 | 20000 | 1772 | 100 | 1173 | 100 | 1014 | 100 | 36 | 99.73 | 20 | 99.45 | 17 | 99.59 |
| 16 | 24000 | 2183 | 100 | 1451 | 100 | 1231 | 100 | 43 | 99.73 | 23 | 99.45 | 21 | 99.59 |
| 19 | 28000 | 2643 | 100 | 1775 | 100 | 1498 | 100 | 56 | 99.86 | 29 | 99.73 | 25 | 99.45 |

<sup>a</sup>The numbers represent total number of NDV reads and maximum 200 reads (optional cut-off value) were used to generate full length consensus sequence. Minimum 5 reads were used as a cut-off to build consensus sequence.
<sup>b</sup>Consensus sequence identity to the reference sequence of NDV LaSota sequenced with MiSeq (MH392212/chicken/USA/LaSota/1946)

Consensus sequences were BLAST searched against NDV custom database
**Note:** Only 28,000 out of total 60,000 reads were utilized for the analysis

### PCR specificity and range of reactivity for NDV genotypes

To determine the utility of the primers for the currently circulating NDV genotypes and the potential cross-reactivity for other AAvVs, which are relatively nonpathogenic in poultry but can confound diagnosis of NDV (45), total RNA from 43 AAvVs, including 23 unique AAvV-1 lineages, (15 different NDV genotypes, 8 different subgenotypes, total samples = 33), as well as AAvV-2–10 and −13 (n = 10) were tested. All AAvV-1 genotypes that are currently circulating globally were amplified with tailed primers; samples 19 and 36 had weak bands of the desired molecular weight compared to other lanes; samples 19, 20, 21, 31 and 32 had an additional, heavier band (~1000 bp). All non-AAvV-1 viruses were negative (Figure S1).

### Quality metrics

The Nanopore QC tool was used to obtain quality metrics plots of all sequencing runs. For MinION runs 3 (6 multiplexed samples) and 4 (33 multiplexed samples), more than 70% of total reads had a quality score greater than ten (Q10 score = 90% accuracy) (Figure S2 A and B). The average overall mean read quality scores in both runs were comparable (run 3 = 10.7, run 4 = 11.0), and the mean quality scores of reads ≥ 10 (mean Q_≥10_) were similar (11.8) for both runs (Figure S2). In addition, analysis of five consecutive batches of reads (each batch = 20,000 reads) obtained at different time intervals from run 4 indicated that the overall mean read quality for each 20,000 read batch remained above 10 (Table S4). Similarly, the mean Q≥10 over time remained consistent in the clinical sample runs (runs 5–7), which had long (12 hrs) sequencing runs (Figure S3, green lines).

### Sub-genotypic resolution of AAvV-1 viruses with MinION sequencing

To determine the capability to effectively detect and differentiate viruses of different genotypes, PCR amplicons from 33 egg-gown isolates, which were representative of 23 different NDV lineages including 15 different genotypes, were barcoded, pooled, and sequenced in a single 12-hour MinION run (run = 4) generating a total of 2.076 million reads. The first 100,000 reads, which were obtained in 3 hours and 10 minutes, were analyzed for identification of all 33 NDV isolates used in the study. All 33 NDV isolates were correctly identified to the sub-genotype level (Table 3). These consensus sequences had 97.82–100% sequence identity to the expected sequence in each of the samples.

**Table 3.**
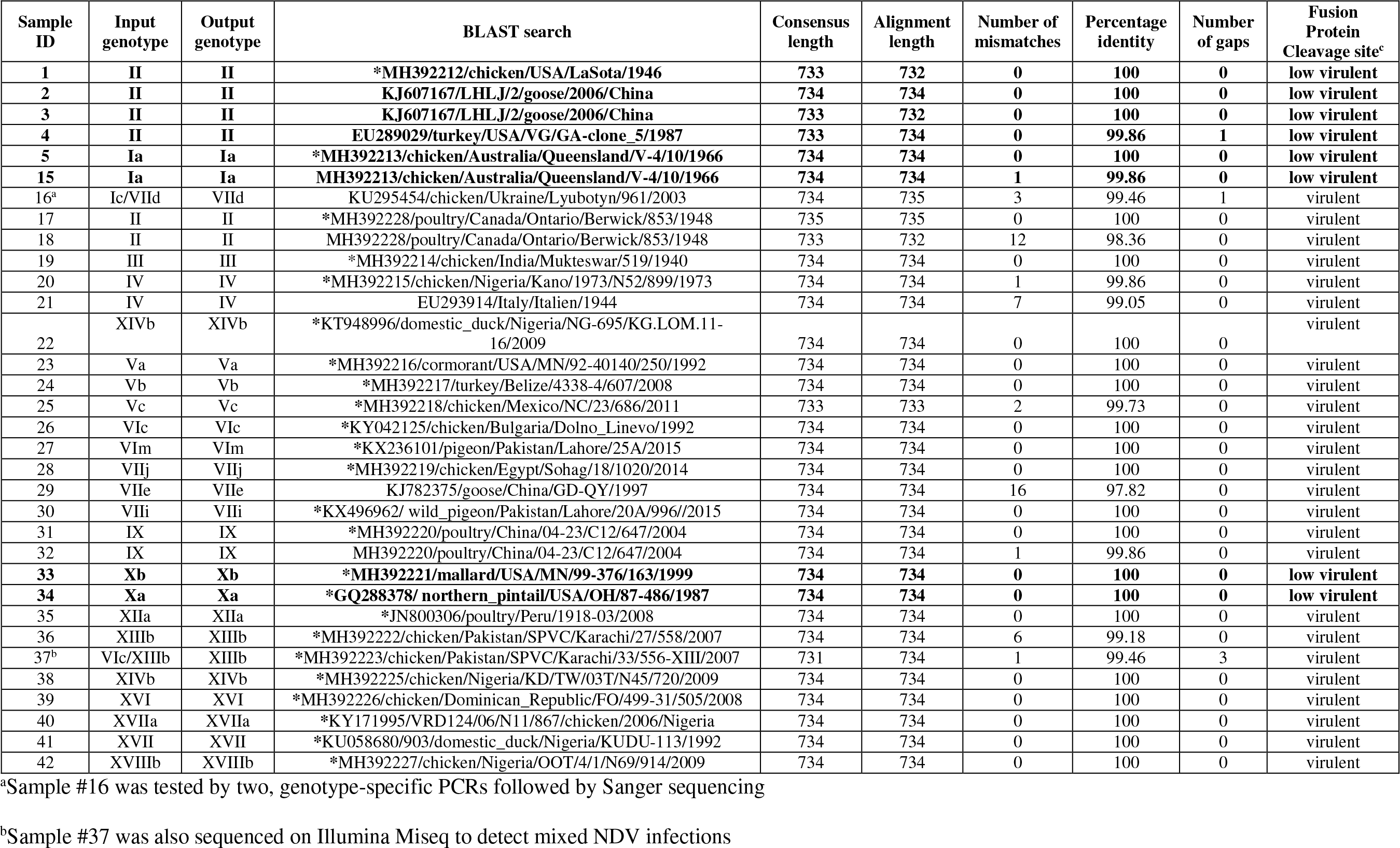
Detection of NDV and consensus sequence identities of pooled 33 NDV genotypes sequenced with MinION and comparison to 23 MiSeq sequences

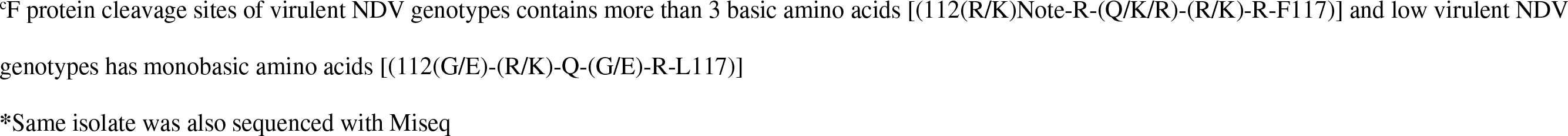

**Note:** Isolates known to have low virulence are highlighted in bold.

While 788 bp was the expected amplicon size, genotypes III, IV, and IX (all previously untested genotypes with this primer set) yielded an unexpected electrophoresis product of ~1000 bp (see above). The sequences obtained from these NDV isolates revealed that in addition to the 788 bp product, an upstream region of NDV genome was amplified, resulting in 1050 bp consensus sequences.

Two virus isolates contained two NDV genotypes. For sample #37, MiSeq detected genotypes XIIIb and VIc, but the AmpSeq workflow only detected genotype XIIIb. Sample #16 had been previously confirmed to contain two genotypes, as determined by two genotype-specific PCRs and Sanger sequencing (data not shown); however, the MinION-based AmpSeq workflow only detected genotype VIId.

### Clinical swab samples from chicken

To assess the potential utility of the protocol on clinical field samples, MinION libraries were generated directly from clinical swab samples. Also, for confirmation, these swab samples were cultured in eggs and the allantoic fluid was sequenced using a MiSeq-based workflow. Out of 11 NDV-positive samples with the culture-MiSeq method, 10 samples were NDV positive by the MinION protocol (Sample #52 being the exception) (Table 4). In the six NDV-positive samples that contained one NDV genotype, as detected by the culture-MiSeq method, the same NDV genotype was also detected with the MinION protocol. The culture-MiSeq method detected two genotypes in samples #45, #46, #47, and #49; whereas, the MinION protocol only detected dual genotypes in samples #45 and #46. In sample #48, only one NDV genotype was detected by culture-MiSeq but two NDV genotypes were detected by the MinION protocol. All 4 samples negative by culture-MiSeq were also negative by the MinION protocol.

**Table 4.**
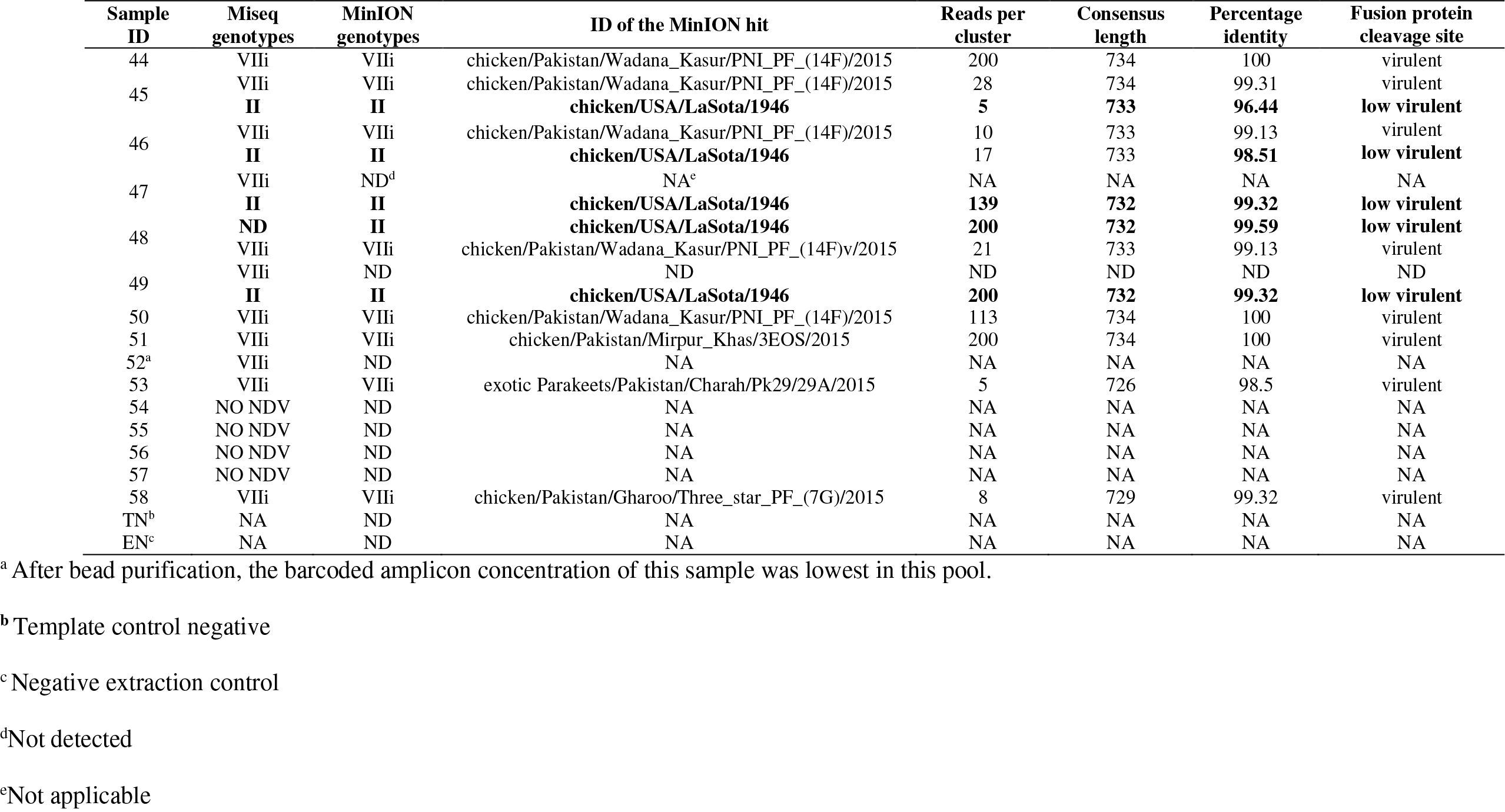
Identification and virulence prediction of NDV genotypes in clinical swab samples collected during disease outbreaks in 2015.

**Note:** Isolates known to have low virulence are highlighted in bold.

### Pairwise comparison of replicated MinION sequences and MiSeq sequences

Pairwise nucleotide distance analysis was used to compare the consensus sequences in six samples across two separate MinION runs, and to compare the MinION consensus sequence to the MiSeq consensus sequence in twenty-four isolates (one isolate representing each genotypes and sub-genotype). There was no variation in the consensus sequence between the MinION runs across those six samples. The MinION and MiSeq consensus sequences were 100% identical, except for sample #37 where the sequence identity was 99.99% to the paired MiSeq sequence. Collectively, these results demonstrate the repeatable accuracy of the MinION-AmpSeq method.

### Phylogeny of NDV genotypes

To confirm the ability of the MinION-acquired partial matrix and fusion gene sequences to be used for accurate analysis of evolutionary relatedness, phylogenetic analysis using consensus sequences (734 bp) obtained from two independent MinION runs (run 3 and 4) was performed. Additionally, the 24 sequences from MiSeq were also included in the phylogenetic tree (Figure 2) to further illustrate the agreement between these two sequencing methods. In the phylogenetic tree, the isolates (n = 33; green font) grouped together with the viruses that showed highest nucleotide sequence identity to them, including those in which MiSeq sequences were available (red font). The six isolates that were sequenced twice (blue font) clustered together. Taken together, the results demonstrated that all sequences clustered to the expected genotype/sub-genotype branch of the phylogram.

**Figure 2.**
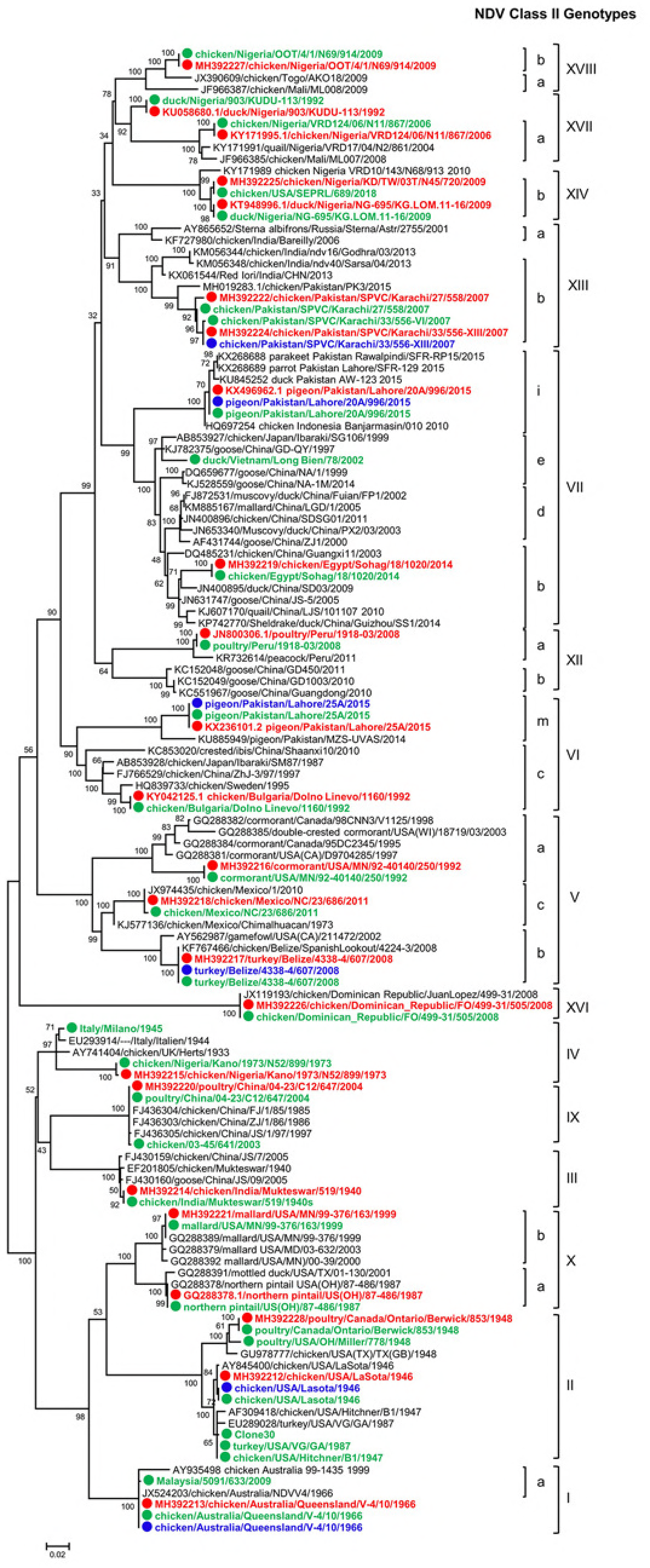
Phylogenetic tree constructed by using the nucleotide sequence (734 bp) of NDV isolates sequenced with MinION, MiSeq and related NDV genotypes from GenBank. The evolutionary histories were inferred by using the maximum-likelihood method based on General Time Reversible model with 500 bootstrap replicates as implemented in MEGA 6. The tree with the highest log likelihood (−9301.9592) is shown. A discrete Gamma distribution was used to model evolutionary rate differences among sites (5 categories [+G, parameter = 0.9325]). The percentages of trees in which the associated sequences clustered together are shown below the branches. The tree is drawn to scale, with branch lengths measured in the number of substitutions per site. The analyses involved 128 nucleotide sequences with a total of 726 positions in the final datasets. The sequences obtained in the current study are denoted with solid circles in front of the taxa name and bold font. Blue circles indicate isolates from MinION sequencing run 1, green circles indicate isolates from MinION sequencing run 2 and red circles indicate MiSeq sequencing.

### Time and cost estimation

The time of sample processing and cost estimation of reagents to multiplex and sequence samples (n = 6; n = 33) from RNA extraction to obtain final consensus sequences is presented in Table S5. From RNA extraction to final consensus sequence calculations, the average time (including sequencing time) to process six samples was approximately 9–10 person-hours and for 33 samples approximately 26 person-hours. Assuming that flow cell can be used multiple times (twice when 33 samples pooled and five times when six samples pooled to prepare one cDNA library) for sequencing, cost per sequencing run and cost per sample were calculated. The cost per sample decreased from $53 (six samples multiplexed) to $31 (33 samples multiplexed).

## Discussion

This study describes a sequencing protocol for rapid and accurate detection, virulence determination, and preliminary genotype identification (with sub-genotype resolution) of NDV utilizing the low-cost MinION sequencer. Additionally, an assembly-based sequence analysis workflow is described for analyzing MinION amplicon sequencing data. This MinION AmpSeq workflow was successful when using a lab repository of egg-grown viruses of varying genotypes, clinical swab samples, and samples containing mixed genotypes.

The sequence heterogeneity among AAvV-1 genomes is well known (3, 46, 47), which complicates the ability to develop a single test that sensitively detects NDV, while also predicting the genotypic classification and virulence. Currently, an RT-qPCR targeting the M gene (9) is most sensitive and is used for screening samples, but this assay only provides positive and negative results of the samples. An RT-qPCR that predicts virulence based on the fusion gene is available (9); however, the lower sensitivity of this assay and the inability of this assay to detect all genotypes (e.g., genotypes Va and VI) (9, 10, 48) complicate diagnostic interpretation when the matrix and fusion tests have conflicting results. Thus, the only truly reliable option to detect some viruses is to design genotype-specific primers and probes (6, 48, 49). For example, Miller *et al* reported that the primer set used in this study detected Class I and all nine of the tested class II genotypes (34); however, this primer set was not tested against other currently circulating genotypes, nor was the sensitivity well defined. The current study includes six additional genotypes, collectively representing all currently circulating genotypes (excluding the Madagascar-limited genotype XI). Additionally, while the preliminary analytical sensitivity of this protocol was determined using only one NDV genotype, the sensitivity of the MinION AmpSeq was comparable to the matrix RT-qPCR and presents an opportunity to implement detection, genotype prediction, and virulence prediction into a single test. As an example of this, in several clinical samples the MinION AmpSeq method detected and identified the LaSota vaccine strain, which is an extensively used vaccine strain for commercial poultry in this region (50). Whereas with the matrix RT-qPCR, these samples would have only been reported as NDV positive and would have required additional testing to determine that the NDV was not virulent.

While the multifaceted nature of this MinION AmpSeq protocol is an advantage, cost and time efficiency must be maintained for it to be useful outside of research laboratories. MinION is inherently rapid due to the real-time nature of the sequencing. Additionally, samples can be multiplexed into a single sequencing run, which reduces time and cost (51). Recently, multiplexing and MinION sequencing of the PCR products from a panel of 5 samples was reported (51). Here a panel of 33 samples was multiplexed while maintaining successful NDV genotyping from data collected within 3 hrs and 10 minutes of sequencing and without affecting mean read quality and percentage of high quality reads.

While Nanopore sequencing has numerous benefits useful for diagnostic testing, the high error rate poses unique challenges to data analysis. Thus, it is important to extract accurate consensus sequences from raw sequencing data (52). As previously discussed, pathogen typing from sequencing data can be done with read count-based profiling or *de novo* assembly approaches (27). However, there are a limited number of available tools suitable for handling the noisy reads currently produced by the MinION platform. The approach in this study takes advantage of the fact that single MinION reads often represent full-length amplicon sequences. By clustering full-length reads based on pairwise identity and subsequently performing consensus calling using standard multiple alignment software, this method quickly and reliably generates accurate *do novo* assemblies from amplicon datasets. Additional refinement using the Nanopolish finisher leads to reliable consensus accuracy >99% with as few as twenty reads per amplicon, and correct genotypic assignments with as few as five reads per amplicon. Thus, this method overcomes the inherently high error rate of Nanopore sequencing and sequence identification and differentiation at the sub-genotype level can be highly reliable.

Because this protocol relies on identity-based clustering prior to assembly, it maintains the ability to detect samples with mixed NDV genotypes, similar to read-based analytical methods. For example, in this study four clinical samples had two different genotypes as detected by MiSeq analysis, two of which were correctly identified by the MinION AmpSeq workflow. In a fifth case, a mixed sample was detected by MinION AmpSeq, but not by MiSeq. A potential explanation for these differences could be that the MiSeq sequencing was performed on egg-amplified samples, which may have altered the relative levels of the two genotypes, as compared to the direct clinical swab sample used for MinION sequencing. Additionally, the differences in molecular techniques (i.e., MinION: targeted; MiSeq: random) may have altered the relative abundance of the genotypes within the sequencing libraries. While further studies into the ability of this workflow to detect and differentiate NDV in samples with more than one genotype are ongoing, rapid NDV genotyping from clinical samples without culturing the virus in SPF eggs has the potential to facilitate disease diagnostics.

Taken together, this protocol reliably detected, genotyped, and predicted the virulence of NDV. A MinION AmpSeq-based approach will be beneficial worldwide and especially in developing countries where the endemicity of high-consequence diseases, such as NDV, and lack of resources are additional challenges to monitoring and studying infectious diseases (50). Furthermore, the advantages of MinION AmpSeq allow for further optimization not possible with other techniques. For example, PCR product length is less of a restriction with MinION AmpSeq as compared to RT-qPCR; thus, there will be one less restriction on primer site design when trying to create a pan-NDV primer set. Work is in progress to utilize the ability of MinION to sequence longer amplicon fragments, which will provide more complete phylogenetic information and may improve the sensitivity of this assay. Overall, MinION AmpSeq improves the depth of information obtained from PCRs and allows for more flexibility in assay design, which can be broadly applied to the detection and characterization of numerous infectious agents.

## Acknowledgments

We gratefully acknowledge Tim Olivier for technical assistance. Special thanks to the Fulbright U.S. Student Program for sponsoring Dr. Salman’s PhD research work. This project was supported by USDA CRIS 6040-32000-072 and DTRA grant 58.0210.5.006 and partially funded by BAA# FRCALL 12-6-2-0015.

## References

1. Miller PJ, Koch G. 2013. Newcastle disease. Diseases of Poultry, 13th ed(Swayne, DE, Glisson, JR, McDougald, LR, Nolan, LK, Suarez, DL and Nair, VL eds), John Wilkey and Sons, Inc, Ames:89–107.

2. Amarasinghe GK, Ceballos NGA, Banyard AC, Basler CF, Bavari S, Bennett AJ, Blasdell KR, Briese T, Bukreyev A, Cai Y. 2018. Taxonomy of the order Mononegavirales: update 2018. Arch Virol:1–12.

3. Dimitrov KM, Ramey AM, Qiu X, Bahl J, Afonso CL. 2016. Temporal, geographic, and host distribution of avian paramyxovirus 1 (Newcastle disease virus). Infect Genet Evol 39:22–34.

4. Commission IOoEBS, Committee IOoEI. 2008. Manual of diagnostic tests and vaccines for terrestrial animals: mammals, birds and bees, vol 2. Office international des épizooties.

5. Aldous E, Mynn J, Banks J, Alexander D. 2003. A molecular epidemiological study of avian paramyxovirus type 1 (Newcastle disease virus) isolates by phylogenetic analysis of a partial nucleotide sequence of the fusion protein gene. Avian Pathol 32:237–255.

6. Kim LM, King DJ, Guzman H, Tesh RB, da Rosa APT, Bueno R, Dennett JA, Afonso CL. 2008. Biological and phylogenetic characterization of pigeon paramyxovirus serotype 1 circulating in wild North American pigeons and doves. J Clin Microbiol 46:3303–3310.

7. Diel DG, da Silva LH, Liu H, Wang Z, Miller PJ, Afonso CL. 2012. Genetic diversity of avian paramyxovirus type 1: proposal for a unified nomenclature and classification system of Newcastle disease virus genotypes. Infect Genet Evol 12:1770–1779.

8. Kim LM, King DJ, Suarez DL, Wong CW, Afonso CL. 2007. Characterization of class I Newcastle disease virus isolates from Hong Kong live bird markets and detection using real-time reverse transcription-PCR. J Clin Microbiol 45:1310–1314.

9. Wise MG, Suarez DL, Seal BS, Pedersen JC, Senne DA, King DJ, Kapczynski DR, Spackman E. 2004. Development of a real-time reverse-transcription PCR for detection of newcastle disease virus RNA in clinical samples. J Clinl Microbiol 42:329–338.

10. Kim LM, Suarez DL, Afonso CL. 2008. Detection of a broad range of class I and II Newcastle disease viruses using a multiplex real-time reverse transcription polymerase chain reaction assay. J Vet Diagn Invest 20:414–425.

11. Fuller CM, Brodd L, Irvine RM, Alexander DJ, Aldous EW. 2010. Development of an L gene real-time reverse-transcription PCR assay for the detection of avian paramyxovirus type 1 RNA in clinical samples. Arch Virol 155:817–23.

12. Ambardar S, Gupta R, Trakroo D, Lal R, Vakhlu J. 2016. High Throughput Sequencing: An Overview of Sequencing Chemistry. Indian J Microbiol 56:394–404.

13. Rhoads A, Au KF. 2015. PacBio sequencing and its applications. Genomics Proteomics Bioinformatics 13:278–289.

14. Chiu CY. 2013. Viral pathogen discovery. Curr Opin Microbiol 16:468–478.

15. Dimitrov KM, Sharma P, Volkening JD, Goraichuk IV, Wajid A, Rehmani SF, Basharat A, Shittu I, Joannis TM, Miller PJ, Afonso CL. 2017. A robust and cost-effective approach to sequence and analyze complete genomes of small RNA viruses. Virol J 14:72.

16. Cruz-Rivera M, Forbi JC, Yamasaki L, Vazquez-Chacon CA, Martinez-Guarneros A, Carpio-Pedroza JC, Escobar-Gutiérrez A, Ruiz-Tovar K, Fonseca-Coronado S, Vaughan G. 2013. Molecular epidemiology of viral diseases in the era of next generation sequencing. J Clin Virol 57:378–380.

17. Marston DA, McElhinney LM, Ellis RJ, Horton DL, Wise EL, Leech SL, David D, de Lamballerie X, Fooks AR. 2013. Next generation sequencing of viral RNA genomes. BMC Genomics 14:444.

18. Gullapalli RR, Desai KV, Santana-Santos L, Kant JA, Becich MJ. 2012. Next generation sequencing in clinical medicine: Challenges and lessons for pathology and biomedical informatics. J Pathol Inform 3:40.

19. Phan H, Stoesser N, Maciuca I, Toma F, Szekely E, Flonta M, Hubbard A, Pankhurst L, Do T, Peto T. 2017. Illumina short-read and MinION long-read whole genome sequencing to characterise the molecular epidemiology of an NDM-1-Serratia marcescens outbreak in Romania. J Antimicrob Chemother 73 (3) 672–679.

20. Greninger AL, Naccache SN, Federman S, Yu G, Mbala P, Bres V, Stryke D, Bouquet J, Somasekar S, Linnen JM. 2015. Rapid metagenomic identification of viral pathogens in clinical samples by realtime nanopore sequencing analysis. Genome Med 7:99.

21. Kilianski A, Haas JL, Corriveau EJ, Liem AT, Willis KL, Kadavy DR, Rosenzweig CN, Minot SS. 2015. Bacterial and viral identification and differentiation by amplicon sequencing on the MinION nanopore sequencer. Gigascience 4:12.

22. Ashton PM, Nair S, Dallman T, Rubino S, Rabsch W, Mwaigwisya S, Wain J, O’grady J. 2015. MinION nanopore sequencing identifies the position and structure of a bacterial antibiotic resistance island. Nat Biotechnol 33:296.

23. Lemon JK, Khil PP, Frank KM, Dekker JP. 2017. Rapid nanopore sequencing of plasmids and resistance gene detection in clinical isolates. J Clin Microbiol 55:3530–3543.

24. Wang J, Moore NE, Deng Y-M, Eccles DA, Hall RJ. 2015. MinION nanopore sequencing of an influenza genome. Front Microbiol 6:766.

25. Quick J, Loman NJ, Duraffour S, Simpson JT, Severi E, Cowley L, Bore JA, Koundouno R, Dudas G, Mikhail A. 2016. Real-time, portable genome sequencing for Ebola surveillance. Nature 530:228.

26. Quick J, Grubaugh ND, Pullan ST, Claro IM, Smith AD, Gangavarapu K, Oliveira G, Robles-Sikisaka R, Rogers TF, Beutler NA. 2017. Multiplex PCR method for MinION and Illumina sequencing of Zika and other virus genomes directly from clinical samples. Nat Protoc 12:1261.

27. Kim D, Song L, Breitwieser FP, Salzberg SL. 2016. Centrifuge: rapid and sensitive classification of metagenomic sequences. Genome Res 26:1721–1729.

28. Caporaso JG, Kuczynski J, Stombaugh J, Bittinger K, Bushman FD, Costello EK, Fierer N, Pena AG, Goodrich JK, Gordon JI. 2010. QIIME allows analysis of high-throughput community sequencing data. Nat Methods 7:335.

29. Schloss PD, Westcott SL, Ryabin T, Hall JR, Hartmann M, Hollister EB, Lesniewski RA, Oakley BB, Parks DH, Robinson CJ. 2009. Introducing mothur: open-source, platform-independent, community-supported software for describing and comparing microbial communities. Appl Environ Microbiol 75:7537–7541.

30. Ip CL, Loose M, Tyson JR, deCesare M, Brown BL, Jain M, Leggett RM, Eccles DA, Zalunin V, Urban JM. 2015. MinION Analysis and Reference Consortium: Phase 1 data release and analysis. F1000Research 4.

31. Schloss PD, Jenior ML, Koumpouras CC, Westcott SL, Highlander SK. 2016. Sequencing 16S rRNA gene fragments using the PacBio SMRT DNA sequencing system. PeerJ 4:e1869.

32. Li C, Chng KR, Boey EJH, Ng AHQ, Wilm A, Nagarajan N. 2016. INC-Seq: accurate single molecule reads using nanopore sequencing. GigaScience 5:34.

33. Alexander D, Swayne D. 1998. Newcastle disease virus and other avian paramyxoviruses, p 156–163. A laboratory manual for the isolation and identification of avian pathogens 4.

34. Miller PJ, Dimitrov KM, Williams-Coplin D, Peterson MP, Pantin-Jackwood MJ, Swayne DE, Suarez DL, Afonso CL. 2015. International biological engagement programs facilitate Newcastle disease epidemiological studies. Front Public Health 3:235.

35. Afgan E, Baker D, Van den Beek M, Blankenberg D, Bouvier D, Čech M, Chilton J, Clements D, Coraor N, Eberhard C. 2016. The Galaxy platform for accessible, reproducible and collaborative biomedical analyses: 2016 update. Nucleic acids research 44:W3–W10.

36. Myers G. Efficient Local Alignment Discovery amongst Noisy Long Reads, p 52–67. In (ed), Springer Berlin Heidelberg,

37. Katoh K, Misawa K, Kuma Ki, Miyata T. 2002. MAFFT: a novel method for rapid multiple sequence alignment based on fast Fourier transform. Nucleic Acids Res 30:3059–3066.

38. Li H, Durbin R. 2009. Fast and accurate short read alignment with Burrows-Wheeler transform. Bioinformatics 25:1754–1760.

39. Li H. 2013. Aligning sequence reads, clone sequences and assembly contigs with BWA-MEM. arXiv preprint arXiv:13033997.

40. He Y, Taylor TL, Dimitrov KM, Butt SL, Stanton JB, Goraichuk IV, Fenton H, Poulson R, Zhang J, Brown CC. 2018. Whole-genome sequencing of genotype VI Newcastle disease viruses from formalin-fixed paraffin-embedded tissues from wild pigeons reveals continuous evolution and previously unrecognized genetic diversity in the US. Virol J 15:9.

41. Thompson JD, Higgins DG, Gibson TJ. 1994. CLUSTAL W: improving the sensitivity of progressive multiple sequence alignment through sequence weighting, position-specific gap penalties and weight matrix choice. Nucleic Acids Res 22:4673–4680.

42. Tamura K, Stecher G, Peterson D, Filipski A, Kumar S. 2013. MEGAxs6: molecular evolutionary genetics analysis version 6.0. Mol Biol Evol 30:2725–2729.

43. Tamura K, Nei M. 1993. Estimation of the number of nucleotide substitutions in the control region of mitochondrial DNA in humans and chimpanzees. Mol Biol Evol 10:512–26.

44. Tamura K, Nei M, Kumar S. 2004. Prospects for inferring very large phylogenies by using the neighbor-joining method. Proc Natl Acad Sci U S A 101:11030–11035.

45. Nayak B, Dias FM, Kumar S, Paldurai A, Collins PL, Samal SK. 2012. Avian paramyxovirus serotypes 2-9 (APMV-2-9) vary in the ability to induce protective immunity in chickens against challenge with virulent Newcastle disease virus (APMV-1). Vaccine 30:2220–2227.

46. Seal BS, King DJ, Bennett JD. 1995. Characterization of Newcastle disease virus isolates by reverse transcription PCR coupled to direct nucleotide sequencing and development of sequence database for pathotype prediction and molecular epidemiological analysis. J Clin Microbiol 33:2624–2630.

47. Seal BS, King DJ, Locke DP, Senne DA, Jackwood MW. 1998. Phylogenetic relationships among highly virulent Newcastle disease virus isolates obtained from exotic birds and poultry from 1989 to 1996. J Clin Microbiol 36:1141–1145.

48. Sabra M, Dimitrov KM, Goraichuk IV, Wajid A, Sharma P, Williams-Coplin D, Basharat A, Rehmani SF, Muzyka DV, Miller PJ, Afonso CL. 2017. Phylogenetic assessment reveals continuous evolution and circulation of pigeon-derived virulent avian avulaviruses 1 in Eastern Europe, Asia, and Africa. BMC Vet Res 13:291.

49. Rue CA, Susta L, Brown CC, Pasick JM, Swafford SR, Wolf PC, Killian ML, Pedersen JC, Miller PJ, Afonso CL. 2010. Evolutionary changes affecting rapid identification of 2008 Newcastle disease viruses isolated from double-crested cormorants. J Clin Microbiol 48:2440–2448.

50. Rehmani SF, Wajid A, Bibi T, Nazir B, Mukhtar N, Hussain A, Lone NA, Yaqub T, Afonso CL. 2015. Presence of Virulent Newcastle Disease Virus in Vaccinated Chickens Farms In Pakistan. J Clin Microbiol 53.5:1715–1718.

51. Wei S, Weiss ZR, Williams Z. 2018. Rapid multiplex small DNA sequencing on the MinION nanopore sequencing platform. G3: Genes, Genomes, Genetics:g3. 200087.2018.

52. Li H. 2016. Minimap and miniasm: fast mapping and de novo assembly for noisy long sequences. Bioinformatics 32:2103–2110.

